# Theta mediated dynamics of human hippocampal-neocortical learning systems in memory formation and retrieval

**DOI:** 10.1101/2023.09.20.558688

**Authors:** Sandra Gattas, Myra Sarai Larson, Lilit Mnatsakanyan, Indranil Sen-Gupta, Sumeet Vadera, Lee Swindlehurst, Paul E. Rapp, Jack J. Lin, Michael A. Yassa

## Abstract

Episodic memory arises as a function of dynamic interactions between the hippocampus and the neocortex, yet the mechanisms have remained elusive. Here, using human intracranial recordings during a mnemonic discrimination task, we report that 4-5 Hz (theta) power is differentially recruited during discrimination vs. overgeneralization, and its phase supports hippocampal-neocortical when memories are being formed and correctly retrieved. Interactions were largely bidirectional, with small but significant net directional biases; a hippocampus-to-neocortex bias during acquisition of new information that was subsequently correctly discriminated, and a neocortex-to-hippocampus bias during accurate discrimination of new stimuli from similar previously learned stimuli. The 4-5 Hz rhythm may facilitate the initial stages of information acquisition by neocortex during learning and the recall of stored information from cortex during retrieval. Future work should further probe these dynamics across different types of tasks and stimuli and computational models may need to be expanded accordingly to accommodate these findings.

## Introduction

The formation, storage and retrieval of episodic memories are thought to rely on interactions between two learning systems, the hippocampus and the neocortex^1–3^. According to the Complementary Learning Systems (CLS) framework, the hippocampus is specialized for rapid acquisition of memory patterns by assigning distinct representations to stimuli regardless of their similarity, overcoming the possibility of catastrophic interference (i.e., pattern separation), while the neocortex is an incremental learner which assigns overlapping representations to stimuli to represent their shared structure and generalize to new stimuli based on their similarity to previously experienced stimuli (i.e., pattern completion)^4–6^.

While existing theories and models are agnostic to the exact direction of hippocampal-neocortical interactions and their timing, inherent to their predictions is that bidirectional information transfer between the two systems needs to occur. CLS posits that the cortex necessarily transfers information to the hippocampus enabling the rapid creation of an index during encoding, while the hippocampus “teaches” experiences to the cortex through slow, interleaved learning during offline periods such as sleep. Recent work has provided evidence for these predicted dynamics. For example, a recent study observed a decrease in neocortical 8-20 Hz power preceding and predicting hippocampal 60-80 Hz power increases during encoding and observed the reverse during retrieval in an associative memory task^7^. The investigators speculated that these events are related to directional information flow from the neocortex to the hippocampus during memory formation, and the hippocampus reinstating a neocortical pattern during retrieval (i.e., pattern completion). Similarly, using representational similarity analyses, another recent study reported early reinstatement of item-context associations in the hippocampus followed by reinstatement of item information in lateral temporal cortex^8^. These studies demonstrate a role for temporally ordered events within the hippocampus and the cortex during an associative memory task that are consistent with model predictions.

Recently, computational models have begun to suggest that hippocampal-neocortical dynamics may vary as a function of general familiarity of the experience and whether pre-existing schema may support its rapid acquisition and integration into neocortex^2,9^. There is evidence for rapid acquisition of schema-consistent items in the neocortex in rodents (on the scale of 48 hours)^10,11^. However, what remains unknown is the degree to which the direction and timing is altered during active learning and recall of schema-consistent items in humans. We therefore sought to test whether there is evidence for bidirectional information flow in both encoding and retrieval, whereby in encoding, in addition to the HC to NC flow at encoding, a HC to NC flow may reflect the initiation of neocortical learning of highly familiar items. Similarly, in retrieval, in addition to the HC to NC flow, a NC to HC flow would reflect access of stored content from the NC to active the hippocampal “index”.

The use of associative memory designs in prior human work leaves another key question unanswered. How do interactions between the hippocampus and neocortex enable the process of behaviorally discriminating among similar “lure” stimuli, which is thought to be dependent on pattern separation^12^? It is possible that a previously encoded similar item needs to be recalled and compared to the currently presented item for the discrimination to occur (recall-to-reject). How does this type of mnemonic challenge, lure discrimination, compare with the nature of the exchange during reinstatement?

Finally, a major question concerns how these systems interact. A contender is the theta rhythm, which is thought to play an important role in facilitating communication across brain regions given its longer period that would accommodate conduction velocity constrains over multi-synaptic communication^13–15^. Theta oscillations may temporally order episodic memory representations and allow for reinstatement of memory representations in the cortex^13,16,17^. Given the theorized role for theta oscillations in mediating interregional interactions and accumulating evidence for its role in learning and memory, we focused on 4-5 Hz frequency range, which overlaps with theta, for mediating NC-HC interactions as a means by which network level communication may arise.

In contrast to past work with associative memory tasks and intra-regional events as a proxy for communication, here we use a well-established and widely used pattern separation task^18^ and an information theory metric to estimate inter-regional directional flow^19^ to test the aforementioned bidirectional interactions between the two systems during both initial memory formation and subsequent discrimination of lures from similar items. Using human depth electrode recordings in patients with epilepsy during performance of a hippocampus-dependent pattern separation task, we demonstrate a prominent role for the dynamics of 4-5 Hz oscillations. We show that 4-5 Hz power is differentially recruited during pattern separation compared to pattern completion, and its phase provides a means of an ongoing bidirectional information transfer within the hippocampal-neocortical network that is task-stage (encoding vs. retrieval) and condition (pattern completion vs. separation) specific. While exchanges between the hippocampus and neocortex were largely bidirectional, small, and significant net directional biases were also present during the formation and recall of pattern separated representations. Somewhat surprisingly, during successful discrimination of similar items (i.e., pattern separation) we observe 4-5 Hz-mediated neocortical→hippocampal directional bias. In contrast, during encoding, we observe 4-5 Hz-mediated hippocampal→neocortical directional bias for items later correctly discriminated at retrieval (subsequent discrimination). Our results identify new dynamical systems interactions underlying pattern separation, and more generally extend findings on the interactions underlying memory formation and retrieval in humans. Additionally, they provide a mechanistic basis for the hypothesized roles of the two learning systems in the formation and retrieval of pattern separated representations.

## Results

Local field potentials were recorded from eight subjects implanted with concurrent neocortical (NC) and hippocampal (HC) intracranial electrodes during a hippocampus-dependent pattern separation task (Table 1, Fig. 1a). The task consisted of an incidental encoding and a retrieval phase. During encoding, subjects viewed images presented serially and indicated with a button-press whether the item belonged indoors or outdoors. During retrieval, either repeated images from encoding (repeat), similar but not identical ones (lure), or new images (new) were shown, and subjects indicated ‘new’ or ‘old’ (Fig. 1a). A ‘new’ response to a lure reflected successful discrimination between encoding and retrieval images (lure+, lure correct), and an ‘old’ response indicated failed discrimination (lure-, lure incorrect).

**Table 1.** Study Participants. Participant demographics, clinical notes, and electrode coverage. R: right, L: left.

| Participant Demographics |  |  |  |  | Electrode Coverage |  |  |  |  |  |  |  |  |
| --- | --- | --- | --- | --- | --- | --- | --- | --- | --- | --- | --- | --- | --- |
| Subject | Age | Sex | Handedness | Years of Education | Seizure onset zone | Orbitofrontal Cortex (OFC) | Frontal Cortex (FRO) | Temporal Cortex (TEMP) | Insular Cortex (INS) | Cingulate Cortex (CING) | Entorhinal/Perirhinal Cortex (EC/PRC) | Hippocampal Dentate Gyrus / CA3 | Hippocampus (HC) |
| 1 | 34 | M | A | 14 | Bilateral HC | 4 | 14 | 7 | 1 | 2 | 2 | 3 | 11 |
| 2 | 32 | F | R | 12 | R lateral TEMP | 5 | 5 | 4 | 0 | 4 | 5 | 0 | 5 |
| 3 | 23 | M | R | 12 | R TEMP | 4 | 18 | 12 | 13 | 6 | 1 | 4 | 9 |
| 4 | 50 | F | R | 16 | Bilateral TEMP | 7 | 7 | 11 | 0 | 6 | 7 | 3 | 13 |
| 5 | 27 | M | R | 12 | Bilateral HC | 2 | 13 | 11 | 0 | 9 | 0 | 3 | 13 |
| 6 | 21 | F | R | 12 | L HC | 4 | 11 | 11 | 3 | 1 | 3 | 1 | 6 |
| 7 | 41 | M | R | 13 | Bilateral lateral TEMP | 2 | 4 | 17 | 2 | 4 | 2 | 0 | 6 |
| 8 | 52 | M | R | 16 | R HC | 1 | 10 | 13 | 6 | 3 | 0 | 0 | 4 |
| Total # of *electrodes* |  |  |  |  |  | 29 | 82 | 86 | 25 | 35 | 20 | 14 | 67 |

**Figure 1.**
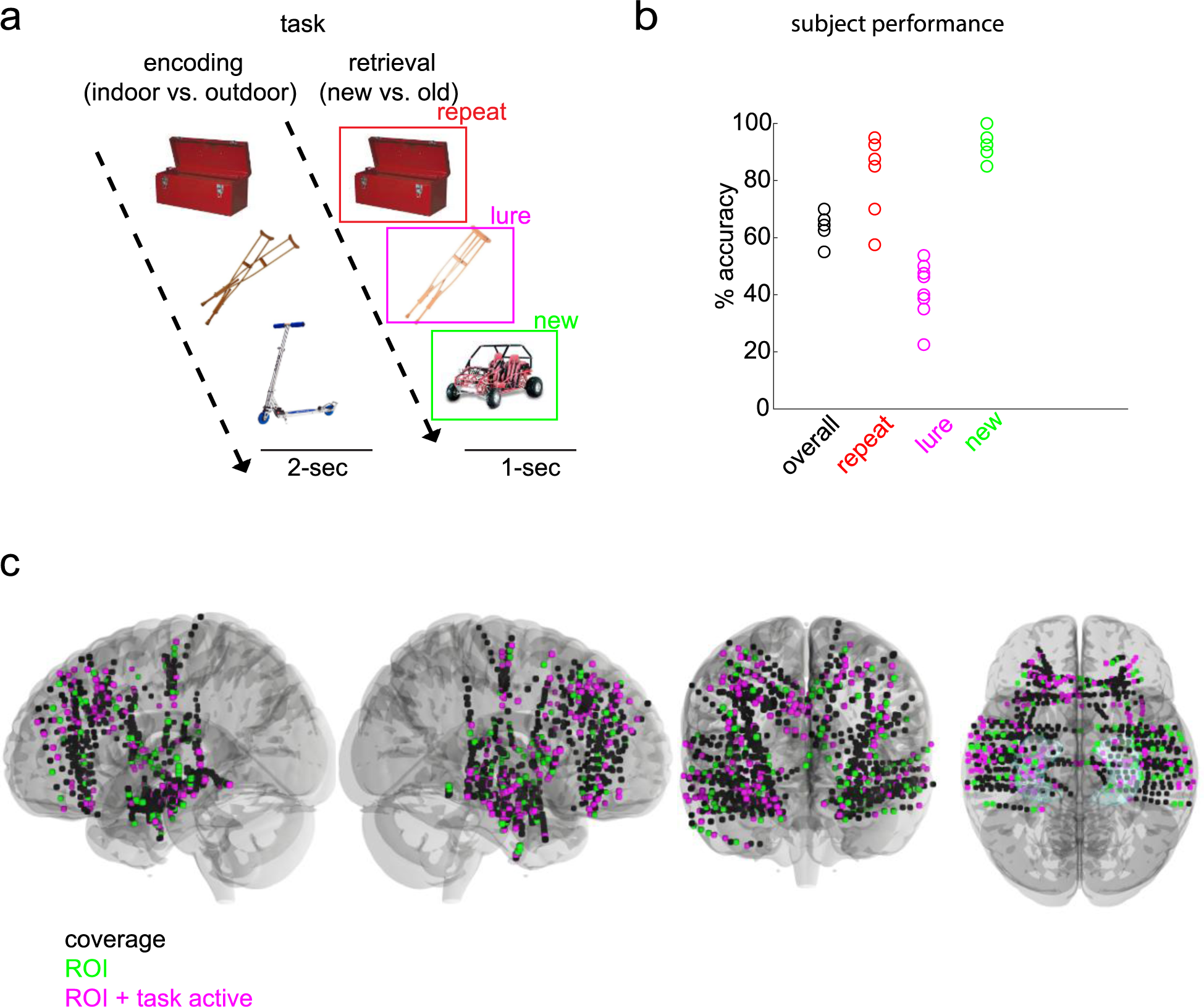
Experimental protocol for investigation of hippocampal and neocortical dynamics. **a,** Task schematic. The task contained two phases—an encoding and a retrieval phase. In the encoding phase, images of objects were displayed one at a time each for a 2-sec duration. Subjects indicated with a button press whether the images belonged indoors or outdoors. In the retrieval phase, one of three image categories was displayed: an image previously shown in the encoding phase (repeat), a similar but not identical image (lure), or a novel image (new). Following a 1-sec image display, and a 0.5 inter-trial period, a question prompt of ‘new or old’ appeared to which subjects responded with a button press. **b,** Individual subject behavioral performance on the task (overall accuracy) and specific task conditions (all, repeat, lures and new images) **c,** Group coverage displayed in standardized (MNI) brain space. Coverage across all subjects (black), neocortical (OFC, FRO, TEMP, CING, INS, EC) and hippocampal ROIs (green), and the task responsive subset of anatomical regions of interest (magenta) are displayed. A hippocampus region is included in the ventral view.

Mean overall task performance was 64.61%, with 82.19% repeat, 92.81% new, and 41.72% lure mean accuracies (Fig. 1b). Neocortical (NC) and hippocampal (HC) cue-responsive electrodes (Table 2, see methods) were utilized for all analyses (Fig. 1c). NC electrodes included orbitofrontal (OFC), lateral and medial frontal (FRO), temporal (TEMP), cingulate (CING), insular (INS), and entorhinal/perirhinal (EC/PRC) cortices. Those for HC included DG/CA3, CA1, and subiculum.

**Table 2.** Proportion of task active electrodes per region during retrieval. Cue responsivity was defined as a significant difference in slow frequency (3-6 Hz) power between pre and post stimulus presentation during retrieval. OFC: orbitofrontal cortex, FRO: frontal cortex, TEMP: temporal cortex, INS: insular cortex, CING: cingulate cortex, EC/PRC: entorhinal/perirhinal cortex, DG/CA3: dentate gyrus / CA3; HC: hippocampus (includes DG/CA3, CA1 and subiculum).

| Subject | OFC | FRO | TEMP | INS | CING | EC/PRC | DG/CA3 | HC |
| --- | --- | --- | --- | --- | --- | --- | --- | --- |
| 1 | 2/4 | 9/14 | 5/7 | 0/1 | 2/2 | 1/2 | 2/3 | 8/11 |
| 2 | 0/5 | 2/5 | 0/4 | 0/0 | 2/4 | 2/5 | 0/0 | 1/5 |
| 3 | 2/4 | 15/18 | 7/12 | 6/13 | 3/6 | 0/1 | 2/4 | 4/9 |
| 4 | 5/7 | 6/7 | 9/11 | 0/0 | 2/6 | 3/7 | 2/3 | 8/13 |
| 5 | 2/2 | 13/13 | 8/11 | 0/0 | 7/9 | 0/0 | 1/3 | 7/13 |
| 6 | 3/4 | 8/11 | 5/11 | 1/3 | 1/1 | 2/3 | 0/1 | 2/6 |
| 7 | 2/2 | 4/4 | 8/17 | 2/2 | 2/4 | 1/2 | 0/0 | 6/6 |
| 8 | 1/1 | 10/10 | 7/13 | 4/6 | 2/3 | 0/0 | 0/0 | 4/4 |

Using serially registered pre- and post-implantation MRI scans, we were able to pinpoint the location of each electrode contact with millimeter precision to achieve subfield-level resolution.

We first identified the neural population level spectral dynamics within the HC that support pattern separation (Fig. 2). Consistent with our prior work^20^ and the well-recognized role for theta in memory processing^13,16,17,21^, ∼4-6 Hz power was significantly higher in the lure+ compared to the lure-condition in the HC (primarily DG/CA3) (*p* < 0.05, paired cluster-based permutation testing) (Fig. 2a). In DG/CA3, only theta survived correction for multiple comparisons (Fig. 2a, bottom panel; see Supplementary Figure S4 for individual entry *p*-values). This was observed on an individual subject basis (Fig. 2c, bottom right panel).

**Figure 2.**
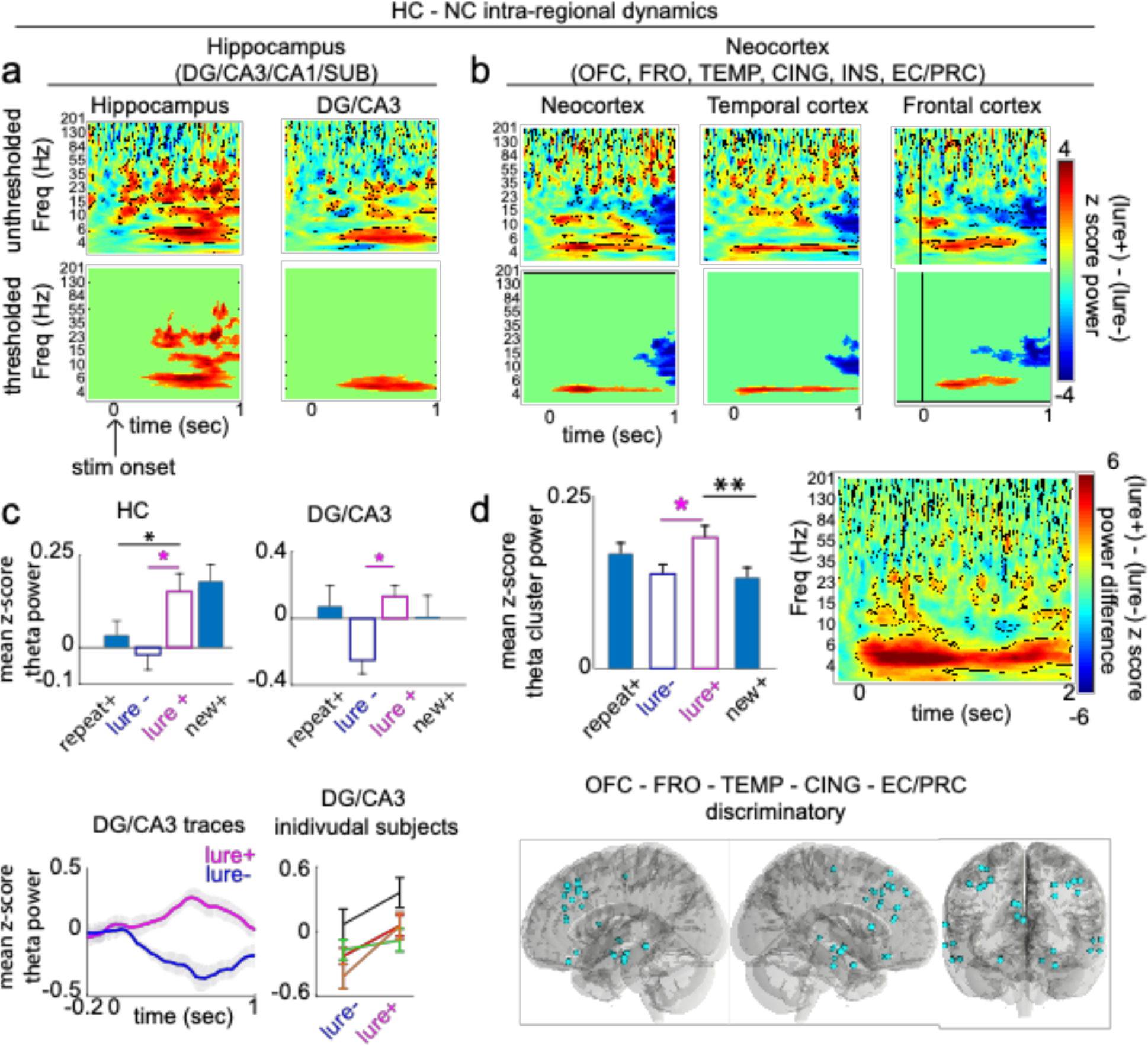
Hippocampal and neocortical intra-regional dynamics during retrieval. **a-b,** z-score difference map (lure+ - lure-) in (a) HC and (b) NC areas. Top: unthresholded maps (outlined voxels, p < 0.05). Bottom: significant clusters after multiple comparisons correction (*p* < 0.05). **c,** Top: mean theta cluster power across 4 condition types (correct (+), incorrect (−)) in HC and DG/CA3. In HC, theta power was significantly higher in the lure+ compared to the repeat+ condition. In DG/CA3, no significant differences between conditions, beyond lure+ and lure-, were observed. Bottom panel: DG/CA3 theta power traces. Individual subject (n=4) DG/CA3 mean theta power. **d,** Top: (left) condition specific theta cluster power in all NC sites. Theta power was significantly higher in the lure+ compared to the new+ condition. (right) Unthresholded difference map using NC discriminatory sites (cyan contacts). Bottom: spatial map of NC discriminatory sites (significantly higher NC theta cluster power in the lure+ compared to the lure-condition; *p* < 0.05). Error shades and bars represent the S.E.M across channels (group analysis) or trials (individual subject and individual channel analysis). \**p* < 0.05, \*\**p* < 0.01, \*\*\**p* < 0.001 indicate significant differences using two-tailed non-parametric paired permutation testing defining channels as observations (1000 permutations).

We then quantified whether this theta rhythm is also recruited in the NC. Across all NC areas, a similar narrow-band, 4-5 Hz power was significantly higher in the lure+ compared to the lure-condition (*p* < 0.05, permutation testing; see Supplementary Figure S4 for individual entry *p*-values) (Fig. 2b). Frontal (∼5-6 Hz) and temporal (∼ 4-5 Hz) sites were key contributors and showed a statistically significant narrow band theta cluster (*p* < 0.05, permutation testing) (Fig. 2b vs. Supplementary Figure 1a). OFC, and to a lesser extent INS sites, also recruited slow-frequency power to a greater extent in the lure+ compared to the lure-condition, however in these areas and in CING and EC/PRC, theta clusters were not significant (Supplementary Figure 1a). A NC theta discriminatory spatial map using the NC cluster is shown in Fig 2d. Theta power was significantly higher in 30 sites across OFC, FRO, TEMP, CING, and EC/PRC in the lure+ compared to the lure-condition (p < 0.05 permutation testing). We next assessed whether theta was the predominant shared rhythm across these discriminatory sites by generating an unthresholded lure+ vs. lure-difference map (Fig. 2d, top right panel). Theta, absent other predominant frequency ranges, was observed. Together, these data suggest that the theta rhythm is recruited by HC and NC during memory processing supporting pattern separation.

To assess specificity and ensure that this differential recruitment of theta was associated with pattern separation above and beyond what might be expected from accurate memory performance, we also compared theta recruitment in lure+ with the new+ and repeat+ conditions (Fig. 2c, d). Hippocampal theta power was significantly higher in the lure+ compared to the repeat+ condition (*p* = 0.0160, permutation testing) but did not significantly differ from the new+ condition (*p* = 0.5754; *p*_(FDR)_ threshold = 0.0160, effect, defined as difference in mean z-score power between groups; see methods, for lure+ vs. repeat+ and lure+ vs. lure-= 0.12 and 0.17, respectively). In DG/CA3 theta power was significantly higher in lure+ compared to lure-(effect = 0.38.) but did not significantly differ from repeat+ nor new+ conditions (lure+ vs. repeat+, *p* = 0.6853, lure+ vs. new+ *p* = 0.2338, *p*_(FDR)_ threshold <0.001; see Supplementary Figure 1c for individual subject data). These patterns in the hippocampus are consistent with our prediction that for lures where pattern separation was hypothesized to occur, the hippocampal response should match that of new+ items, whereas for lures where pattern separation did not occur, the hippocampal response should resemble that of repeat+ items. However, also plausible is that the shared increased theta power for lure+ and new+ is due to outcomes of the underlying memory computations; such outcomes include a novelty signal (could be part of a pattern separation signal), the decision that the item is new, and the behavioral motoric response of new. These are shared processes between the lure+ and new+ and not present in the other conditions. In the NC, theta power was significantly higher in the lure+ compared to the new+ condition (*p =* 0.002) but did not significantly differ from the repeat+ condition (*p* = 0.1319, *p*_(FDR)_ threshold = 0.002); effects for lure+ vs. lure- and lure+ vs. new+ = 0.053 and 0.058, respectively (Fig. 2d). Together, these results support the notion that theta power in the HC and NC was higher for conditions where pattern separation occurred, and power in the NC was particularly higher in the condition where pattern separation was taxed (lure+). Importantly, the observed theta power increases were robust to several referencing techniques including our employed Laplacian referencing (minimizing local commonalities in time domain signal) and distant white matter (minimizing commonalities in time domain signal across distant areas), persisted after subtracting the mean evoked response across all trials in a given channel, and was robust to different normalizations (condition specific peristimulus normalization and normalization relative to the entire recording).

Guided by the notion that the longer period of the slow-frequency theta rhythms may facilitate distant interactions limited by conduction velocity across longer synaptic delays^14^, we next asked whether theta phase provides a means for behaviorally relevant dynamical interactions between the two learning systems. More specifically, we hypothesized that during retrieval, there would be epochs whereby theta mediates NC → HC information transfer, possibly reflecting the access of memory content from NC stored patterns. To do this, we used an information theory metric, phase transfer entropy (PTE) (see methods), to estimate directional 4-5 Hz information transfer. A PTE value of 0 denotes pure bidirectional interactions, > 0 denotes a larger weight to the NC→ HC direction, and < 0 denotes a larger weight to the HC→NC direction. All cue responsive electrodes during the retrieval stage of the task that were in the NC (OFC, FRO, TEMP) or the HC were included for all PTE analyses.

The two systems interacted at retrieval in both directions, with distinct dynamics (Fig. 3a). When collapsing across the ongoing bidirectional nature of the exchange, the net interactions favored the NC→ HC direction during the lure+ compared to the lure-condition (*p* = 0.002, permutation testing, *p*_(FDR)_ threshold = 0.002, effect = 0.004; Fig. 3b; see additional chance statistical analysis in Supplementary Figure S6a). This pattern was primarily driven by prefrontal cortical sites (Supplementary Figure 1e), robust to the choice of time lag between the NC and HC (Supplementary Figure 1d) and was observed on an individual subject basis (7/8 subjects) (Fig. 3c). Investigating dynamical interactions further as a function of time (PTE-FOT) (Fig 3a) revealed sustained significant differences favoring the NC→HC direction in the 0.6-1 sec post-stimulus period. However, present earlier in the post-stimulus period (∼0-0.6 sec) were more frequent alternations between the two directions, including the HC→NC bias. A repeated measures ANOVA revealed a significant main effect for time on NC-HC interactions F (50,73400) =18.407, lower-bound adjusted *p*-value for sphericity *p_LB_*< 0.001, *η*^2^ = 0.012 and a significant interaction between time and condition, *F* (50,73400) = 14.566, *p_LB_* < 0.001, *η*^2^ = 0.010. Clear epochs with significant differences between the lure+ and lure-conditions were subsequently identified with FDR corrected permutation testing (*p*_(FDR)_ threshold = 0.019).

**Figure 3.**
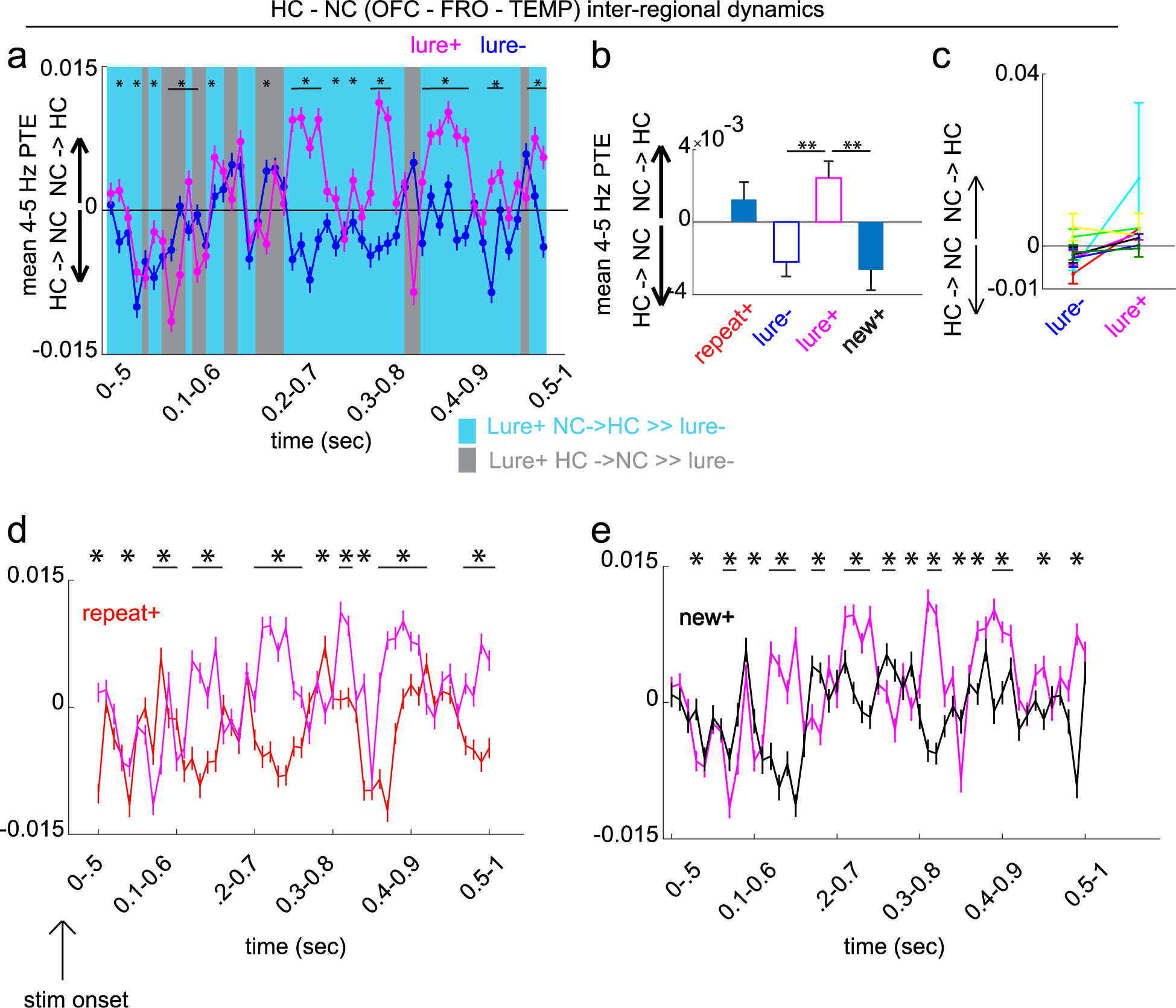
Hippocampal and neocortical inter-regional dynamics during retrieval. **a,d,e,** Phase transfer entropy (PTE) as a function of time (FOT) during the 1-sec post-stimulus period contrasting lure+ with (a) lure-, (d) repeat+ and (e) new+. Blue shading indicates epochs with larger NC→HC weight for lure+ compared to the contrasted condition, while gray shading indicates epochs with larger HC→NC weight. **a,** Lure+ vs. lure-, repeat+ and new+ PTE FOT main effects for time and condition-time interactions (*p_LB_* < 0.001 for all tests; lower-bound adjusted *p*-values for sphericity). PTE significantly differed in particular time epochs between the lure+ and each of the three conditions. **b,** Same as (a,d,e) but for a single time window (0-1 sec), comparing lure+ with all conditions. NC→HC directional bias was higher in the lure+ compared to the lure- and new+ conditions. **c,** Display of individual subject data for the lure+ vs. lure-group level contrast in (b). Error bars represent the S.E.M across channel pairs (group analysis) or trials (individual subject analysis). \**p* < 0.05, \*\**p* < 0.01, \*\*\**p* < 0.001 indicate significant differences using two-tailed non-parametric paired permutation testing defining channel pairs as observations (1000 permutations).

Different dynamical interactions were observed in the new+ and repeat+ conditions, when examining both the time average and interactions as a function of time. The NC→HC bias across the 1-sec post-stimulus period was also significantly higher in the lure+ compared to the new+ condition (*p* = 0.004, effect = 0.005), but did not significantly differ from the repeat+ condition (*p* = 0.360, *p*_(FDR)_ threshold = 0.002) (Fig 3b). However, distinct theta PTE dynamics were observed when considering the time-resolved measure (Fig 3d-e). When considering the lure+ and repeat+ contrast, a repeated measures ANOVA showed a significant main effect for time on NC-HC interactions F (50,73400) =19.343, *p_LB_*< 0.001, *η*^2^ = 0.013 and a significant interaction between time and condition, *F* (50,73400) = 17.8, *p_LB_* < 0.001, *η*^2^ = 0.012. Epochs that significantly differed were again identified with FDR corrected permutation testing (*p*_(FDR)_ threshold = 0.002), which showed that the condition difference in NC→HC bias occurs earlier compared to the contrast with the lure-condition. Lastly, lure+ time-resolved theta PTE differed from that of new+; a repeated measures ANOVA showed a significant main effect for time on NC-HC interactions F (50,73400) =17.044, *p_LB_* < 0.001, *η*^2^ = 0.011 and a significant interaction between time and condition, *F* (50,73400) = 14.156, *p_LB_* < 0.001, *η*^2^ = 0.010. Like the contrast with the new+, individual time epoch analysis showed that the NC→ HC bias occurred earlier relative to stimulus onset compared to the lure+ vs. lure-contrast (permutation testing, *p*_(FDR)_ threshold = 0.022). Altogether, HC and NC intra-regional theta dynamics estimated with power analysis coupled with inter-regional dynamics estimated with phase analysis differentially support the lure+ condition whereby pattern separation is taxed.

Next, we investigated the role of theta dynamics during encoding of items that were subsequently discriminated correctly during retrieval (i.e., a subsequent memory analysis). First, we tested whether theta recruitment is larger for items later discriminated (→lure+) compared to those associated with failed discrimination (→lure-). During encoding, 4-5 Hz power was recruited for both stimulus types in HC and NC (Fig 4a). In the retrieval task active sites (same sites used in Fig. 2), power was numerically higher in the →lure+ compared to the →lure-condition at a slightly different center frequencies (4 Hz for HC and 3 and 5 Hz for NC; Supplementary Figure 2d), however these clusters are not significant after multiple comparisons correction. In contrast, when limiting NC analysis to the sites discriminating lure+ and lure-trials during retrieval, we observed a statistically significant cluster including the theta range, in which encoding cluster power was significantly higher in items later discriminated (*p* < 0.05, permutation testing) (Supplementary Figure 2c, traces displayed in Fig. 4b). It is important to note that all analyses include HC and NC electrodes that were task active during retrieval—not encoding. Encoding theta power results using this subset of electrodes are marginal. Interestingly, when using all HC electrodes, however, consistent with prior reports regarding hippocampal theta during encoding^13^, observed is a reliable theta cluster (higher center frequency - 10 Hz) that is significantly higher for (→lure+) compared to (→lure-) items (Supplementary Figure S5). This raises a possibility that hippocampal sites are differentially recruited during encoding and retrieval and at different center frequencies for theta.

**Figure 4.**
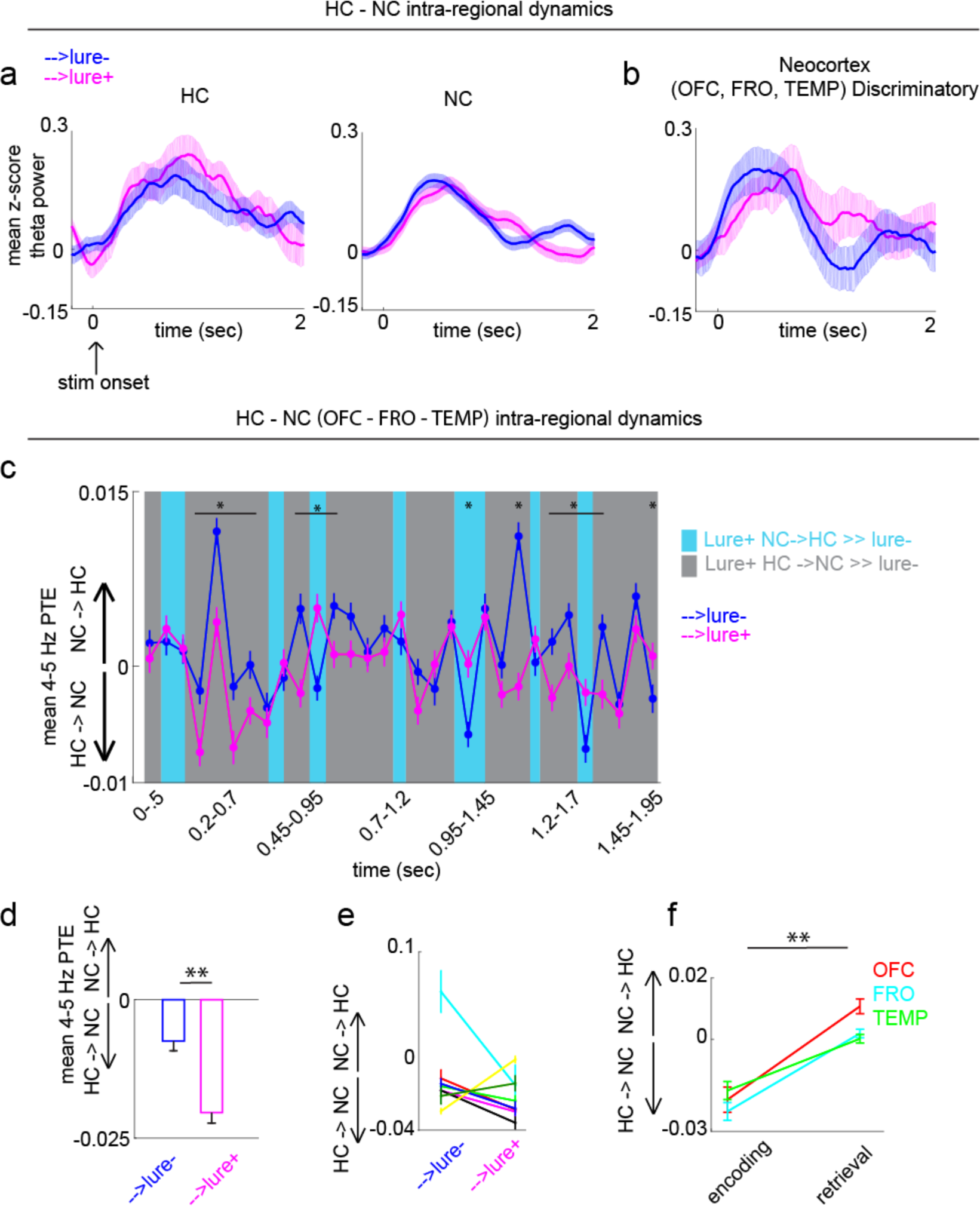
Hippocampal and neocortical intra- and inter-regional dynamics during encoding. **a**, Encoding HC and NC mean 4-5Hz power time traces for stimuli which were later discriminated (→ lure+) and those associated with failed discrimination (→lure-) in retrieval. **b,** Same as in (a), but limiting NC sites to those showing a lure+ vs. lure-discrimination signal during retrieval. During encoding a statistically significant cluster after multiple comparisons correction (Supplementary Figure 2c) including the 4-5 Hz range was observed in these sites. **c,** PTE as a FOT during the 2-sec post-stimulus onset (main effect for time *p_LB_* = 0.016 and condition-time interaction *p_LB_* = 0.010). PTE significantly differed in particular time epochs between the lure+ and lure-conditions. Blue shading indicates epochs with larger NC→HC weight while gray shading indicates epochs with larger HC→NC weight. **d-e,** Same as in (c) but for (d) a single time window (0-2 sec), and (e) in individual subjects. **f,** HC-NC PTE values in the →lure+ encoding, and lure+ retrieval conditions, shown separately in each NC site. Error shades and bars represent the S.E.M across channels (power), channel pairs (PTE), or trials (individual subjects). \**p* < 0.05, \*\**p* < 0.01, \*\*\**p* < 0.001 indicate significant differences using two-tailed non-parametric paired permutation testing defining channel/channel pairs as observations (1000 permutations).

We then asked whether theta-mediated systems interactions during encoding are important for later discrimination. We compared encoding and retrieval PTE values separately in OFC, FRO and TEMP (same contacts used for the retrieval analysis but displayed separately per region) with the HC. In each region pair, the HC→NC directional bias was more prevalent during encoding, while the opposite direction of information flow was more prevalent during retrieval (Fig. 4f) (*p* = 0.002 for each site, permutation testing, *p*_(FDR)_ threshold = 0.002, effects for OFC, FRO and TEMP = 0.0304, 0.0254, and 0.0169, respectively. To test the second hypothesis, we quantified information transfer in the two condition types over the 2-second post-stimulus period in encoding, and as a function of time in the same period. Encoding interactions were again largely bidirectional (centered around 0), with a significantly larger HC →NC information flow in →lure+ compared to →lure-conditions (*p* = 0.002, effect = −0.0129; Fig. 4d; see additional chance statistical analysis in Supplementary Figure S6b). This pattern was robust to the choice of time lag between the NC and HC (Supplementary Figure 2f) and was observed in most individuals (6/8 subjects) (Fig. 4e). PTE-FOT analysis further identified epochs (∼0.2-0.7 sec, and 1-2 sec) where the HC→ NC bias was predominant (Fig. 4c). A repeated measures ANOVA revealed a significant main effect for time on NC-HC interactions F (30, 44040) = 5.861, *p_LB_* = 0.016, *η*^2^ = 0.004 and a significant interaction between time and condition, *F* (30, 44040) = 6.6476, *p_LB_* = 0.010, *η*^2^ = 0.005. Once again, epochs with significant PTE differences between the lure+ and lure-conditions were identified with FDR-corrected permutation testing *(p*_(FDR)_ threshold = 0.02). Collectively, the above results suggest that while systems interactions are largely bidirectional, small biases in directionality during encoding and retrieval may provide a mechanism for the HC’s role as an index and the NC’s role as the storage site for encoded patterns.

Finally, we investigated whether learning system dynamics supporting early mnemonic discriminatory processes are distinct from the dynamics supporting subsequent response generating mechanisms (e.g., executive function and motor planning) associated with such processes. Within a trial, we refer to the early phase consisting of the 1-sec display of an image as the ‘mnemonic epoch’, and the later phase consisting of the last 1-sec preceding the subject response (while the question prompt is on the screen) as the ‘response generating epoch’. These two epochs were non-overlapping in time. In both systems, a drop in power across a broad range of frequencies (∼3-30 Hz) preceding subject response was significantly larger in the lure+ compared to the lure-condition (*p* < 0.05, permutation testing; see Supplementary Figure S4 for individual entry *p*-values) (Fig. 5a-d). In contrast, this was not statistically observed in the NC discriminatory sites (Fig. 5f). DG/CA3 slow-frequency power drop was also significantly larger in the lure+ compared to the new+ (*p* = 0.0240, paired permutation testing), but not the repeat+ condition (*p* = 0.2957, *p*_(FDR)_ threshold = 0.0240; Fig. 5g); effects for lure+ vs. lure- and new+ contrasts = −0.380 and −0.286, respectively. NC slow-frequency power drop was specific to response judgements associated with successful pattern separation (lure+ vs repeat+ *p* = 0.002, lure+ vs. new+ *p* = 0.002, *p*_(FDR)_ threshold = 0.0279, effects for lure+ vs. lure-, new+ and repeat+ contrasts = −0.0492, −0.0749 and −0.0717, respectively; Fig. 5h).

**Figure 5.**
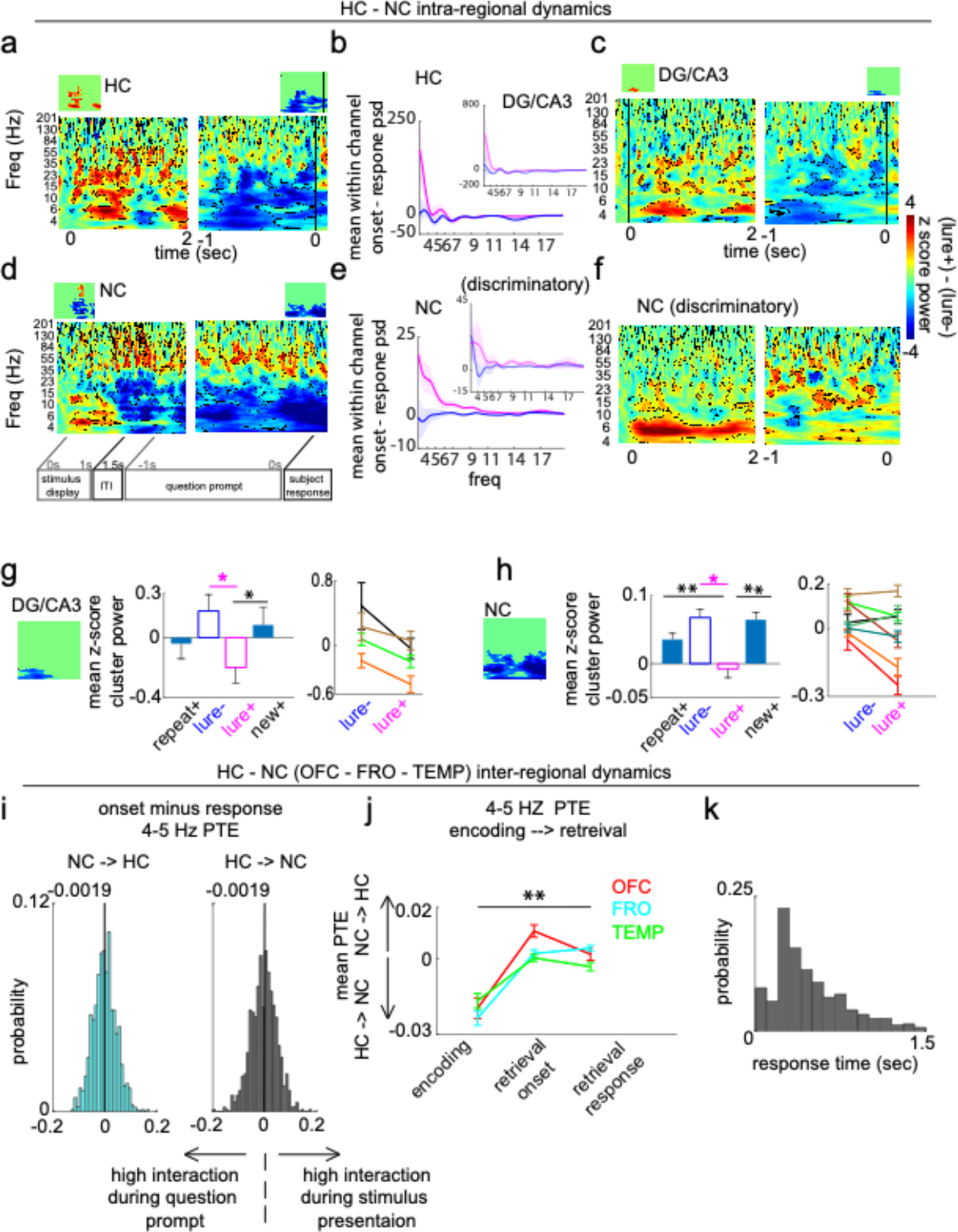
Hippocampal and neocortical intra- and inter-regional dynamics during retrieval response period. **a,c,d,f,** z-score unthresholded and thresholded (insets) difference maps (lure+ - lure-) locked to stimulus onset (left) and response (right) in (a) HC, (c) DG/CA3, (d) NC, and (f) discriminatory NC sites. **b, e,** Mean within-channel subtraction of power spectral density between the 1-sec post stimulus and 1-sec pre-response periods (onset - response PSD) in the (b) HC and NC (e) areas. **g, h,** Mean response locked slow frequency cluster (insets) power in DG/CA3 (g) and NC (h) across 4 conditions on a group, and 2 conditions on an individual subject level. **g,** In DG/CA3, slow frequency power was significantly lower in the lure+ compared to the new+ condition. **h,** In NC, power was significantly lower in the lure+ compared to the repeat+ and new+ conditions. **i,** Probability density function of within channel onset minus response 4-5 Hz PTE in the NC→HC (cyan) and HC → NC (black) directions). Median values are indicated on top of each distribution. **j,** Extension of Fig. 4f to include the response task period; PTE values in the →lure+ encoding, and lure+ retrieval onset and response conditions, shown separately in each NC site. **k,** Response time distribution across trial types and subjects. Error shades and bars represent the S.E.M across channels (power), channel pairs (PTE), or trials (individual subjects). \**p* < 0.05, \*\**p* < 0.01, \*\*\**p* < 0.001 indicate significant differences using two-tailed non-parametric paired permutation testing defining channel/channel pairs as observations (1000 permutations).

We also tested whether this slow-frequency power drop was associated with changes in systems interactions between onset and response periods by estimating the distribution of within-channel onset-minus-response PTE, separately in each direction (Fig. 5i). With equal probability, NC-HC pairs are likely to increase or decrease the degree of information transfer upon transition to the pre-response period. This was observed in NC (OFC-FRO-TEMP) 4-5 Hz (Fig. 5i) and across 3-20 Hz (Supplementary Figure 3h), and across all NC areas irrespective of discrimination level (Supplementary Figure 3i). Amidst these interaction shifts, the net direction during the pre-response period was a larger NC → HC bias compared to that observed in encoding (*p* = 0.002 for each site, paired permutation testing, *p*_(FDR)_ threshold = 0.002; effects for OFC, FRO and TEMP = 0.0213, 0.0275, and 0.0135, respectively; Fig. 5j). Overall, these results suggest that during the later phase of response preparation and generation (presumably after the mnemonic processing has occurred), there is a drop in theta power in DG/CA3 and NC particularly for the lure+ condition, as well as an equal probability of a change in directional flow between the two directions. This analysis demonstrates that power dynamics during the response generating epoch are distinct from the mnemonic epoch, but the net direction of phase mediated information flow is relatively maintained from the mnemonic to the response generating epoch.

## Discussion

In summary, we have shown that 4-5 Hz power in the HC (particularly DG/CA3) and NC supports pattern separation, and that its phase provides a potential mechanism for bidirectional information transfer (measured by phase transfer entropy). Consistent with prior theoretical and experimental work, encoding NC→ HC flow and retrieval HC→ NC were indeed observed. However, somewhat counter-intuitive, is that the *net* directions during encoding was a HC → NC flow and during retrieval was a NC→HC flow. The early increases in 4-5 Hz power during the mnemonic epoch are followed by a pre-response drop; however, information flow distributions appear to be largely sustained. Overall, these results provide a new, expanded account of the nature of bidirectional hippocampal-neocortical information exchange related to encoding and retrieval of episodic memories.

Limitations of prior work investigating NC and HC involvement in humans include difficulties with making inferences about directionality^8^, lack of concurrent HC and NC recordings^22^, extent of neocortical coverage^7^, and behavioral paradigms that do not directly tax pattern separation^7,8,22^. Here, we identify the dynamics of 4-5 Hz spectral power and phase-dependent directional interactions within and between the HC and the NC. Neocortical areas included orbitofrontal, cingulate, and additional frontal sites, and insular, entorhinal perirhinal and additional temporal sites. Dynamics mapped were those supporting memory formation (new+) and retrieval (repeat+), as well as formation and retrieval of events underlying successful pattern separation (lure+) vs. overgeneralization errors (lure-). Each of these cognitive states is supported by unique interactive dynamics between the HC and NC. These results provide empirical evidence for the role of theta and theta-mediated interactions in supporting pattern separation, and more generally memory formation and retrieval. They form a mechanistic basis for the long-hypothesized role of theta in episodic memory processing and expand upon the bidirectional interactions predicted by memory indexing theory^1,23^ and computational models of the two learning systems^2,4–6,9^.

It is critical to note that bidirectional interactions between HC and NC have been recently investigated in human physiology recordings^7^. In this study, the authors report that decreased NC alpha/beta (8-20 Hz) power precedes and predicts increased HC fast gamma (60-80 Hz) during episodic memory encoding, whereas increased HC slow gamma (40-50 Hz) power precedes and predicts decreased NC alpha/beta power during retrieval. While at first our results may appear at odds with these findings, there are key differences across studies. Like their findings, we qualitatively observed decreased NC alpha (∼7-15 Hz) during encoding for subsequently discriminated items (Supplementary Figure S2a, d). We also observed increased HC slow gamma (∼35-55 Hz, Fig 2a, 5a) and decreased NC alpha/beta (∼12-23 Hz, Fig 2b) for successfully discriminated items during retrieval. While these spectral dynamics were shared between the two studies, we chose to focus *a priori* on the theta rhythm given its known role in mnemonic processes and long-range communications. In our results, it also was a common signal across the NC and HC (in particular DG/CA3), which makes it a likely candidate for information transfer. We used an information theoretic approach (phase transfer entropy) to directly estimate information transfer across regions. The latter cannot be inferred purely from correlation analysis. We demonstrate dynamics of bidirectional information exchange, which include the directions reported by the study, however our data suggests a more complex interplay among the two regions during both encoding and retrieval and different net directions. Thus, our reported results complement previously published work^7,8^ and the more salient directions inferred from prior theoretical work^4,23^, but also clarify the richness of such exchange, that it is constantly changing in direction in an encoding and retrieval-dependent manner, and that the *net* directions of such exchanges, at least measured through the phase of 4-5 Hz, differs from the salient directions predicted by prior work.

Since the 1930s, the theta rhythm has been observed across species and behavioral states^16,24^. Amongst a diverse set of its behavioral correlates, a subset includes arousal, attention, decision making, learning and memory, working memory and motivation^16,21^. These processes all require the processing of external inputs^16^. The hippocampal theta rhythm is thought to function as a temporal organizer, enabling first and higher order temporal linkages amongst cell assemblies and converting such orders to synaptic strengths^16^. This is supported by experimental work showing that after controlling for other factors, theta was primarily associated with temporal ordering of experiences^21,25,26^. Theta cycles may separate encoding and retrieval epochs by alternating processing modes during the peak and trough^17^. During the peak, LTP occurs and more predominant inputs from EC to CA1 prevail, possibly facilitating encoding, while LTD occurs during the trough with predominant CA3 inputs to CA1, possibly facilitating retrieval^27,28^. Evidence for theta’s role has been demonstrated in its temporal organization of place^29^ and grid^30^ cell firing (phase precession)^31^, and cells encoding time intervals^32^. These findings point to theta as a potential mechanism to encode space and time, the basic ingredients for episodic memory, thereby serving as a fundamental neural code in memory processing.

With respect to learning systems interactions, theta is proposed to encode and temporally bind representations across neocortical modules, as well as mediate the interactions between HC and NC with the HC acting as an index generator^13^. Our results could be explained by this model, providing a mechanistic basis for memory indexing theory and similar models. The involvement of the two learning systems in this hippocampus-dependent pattern separation task is predicted by theoretical and computational models and is demonstrated by theta recruitment in HC and NC sites during both encoding and retrieval. While these models are largely agnostic to the exact nature of directional information transfer, they do predict bidirectional information transfer between the HC and NC. This is indeed what we observe—that information transfer is largely bidirectional (PTE centered around 0) with fluctuations between the two directions as a function of time (PTE > and < 0). Models further predict that during encoding a unique spatiotemporal neocortical pattern is generated upon an experience, which is strengthened by the hippocampus^23^. This NC pattern – information about the current experience – is likely communicated to the HC during encoding (NC→HC); a direction observed in our data during both the encoding and retrieval phases. A limitation worth noting is the absence of electrode coverage in the ventral visual stream (occipital and ventral temporal cortex), which hinders us from fully exploring how sensory information about current experience is communicated to the hippocampus.

In addition to the salient prediction that NC communicates information about the experience to the HC during encoding, we assessed the possibility that HC → NC communication also occurred. Interestingly, this less-emphasized direction was indeed the predominant directionality we observed during encoding. It is possible that this directional exchange constitutes the initial stages of acquisition of information by the cortex from the hippocampal “teaching signal”, a process previously thought to be dependent on offline periods or repeated experiences but is possibly primed during acquisition. Another possibility is that a unique spatiotemporal code is communicated from hippocampus to the cortex. These two possibilities may in part reflect rapid integration and learning by the cortex. The CLS model was recently updated to predict that the neocortex is capable of rapid learning if new information is consistent with existing schemas^2,9^, which is supported by experimental evidence^10,11^. Given the schema-consistent everyday objects viewed in this study, it is possible that integration of information into neocortical connections occurs rapidly in this task. However, future work is needed to specifically test these possible interpretations.

During retrieval, models propose that upon encountering a partial cue, neocortical modules can be activated and accessed by the hippocampus to facilitate full retrieval. We observe that the increase in NC theta power precedes that of the HC during successful retrieval and bidirectional information flow is biased toward the NC→HC direction. These two observations could reflect reactivation of a neocortical pattern and its access by the hippocampus to enable comparison of the similar lure to an existing memory, i.e., recall-to-reject^5,33^, a likely needed process for accurate discrimination. In line with the notion that 4-5 Hz power likely supports the aforementioned mnemonic processes, is that slow frequency power was significantly diminished preceding response. This, in conjunction with prior work showing increased hippocampal and DLPFC gamma power in the same task preceding response^35^, which we replicated in the present study (Fig 5a, d), suggest relatively more local neocortical and hippocampal activity during the decision phase.

A possible alternative explanation for 4-5 Hz power dynamics during successful encoding and retrieval is active sensing and exploration that facilitates memory encoding^16,24,34–37^. This is a possible explanation for power increases during encoding of items that are subsequently discriminated (Supplementary Figure S2a, c for NC and S5 for HC). However, this is unlikely to be driving condition differences during retrieval since the new+ condition is also novel but does *not* show similar increases in power or interaction dynamics to the lure+ condition. Another possibility is that theta facilitates retrieval of the context in which memory of the item was generated rather than memory of the item itself^38,39^. However, this again would not explain the differences between lure+ and repeat+, both of which presumably have accurate stimulus-specific context retrieval that leads to accurate recall. Finally, an intriguing possibility is that oscillatory inhibition varies as a function of theta phase, with increasing levels of inhibition promoting the suppression of competing memory traces to facilitate accurate retrieval^21,40^. This could be a mechanism by which encoding and retrieval dynamics contribute to successful discrimination among similar items, although this possibility needs to be directly tested by manipulating inhibition.

It is important to distinguish the 4-5 Hz activity observed here from frontal-midline theta (FMT) activity. FMT has been observed in several neocortical areas, including the anterior cingulate, dorsolateral prefrontal cortex and other components of the default mode network^21^. It is thought to be associated with monitoring response conflict for example in Stroop tasks^41^, inhibitory motor control^42,43^ and attention and adaptive control^44^ as well as response conflict during value-based decision tasks^45^. More generally, midfrontal theta has been hypothesized as a mechanism for cognitive control^46^. Since the recall-to-reject processes, hypothesized to be employed in the lure+ condition, relies on some degree of cognitive control and there is evidence that theta waves do propagate throughout the human neocortex^47^, we cannot rule out the possible contributions of midline frontal theta to our findings of 4-5 Hz neocortical dynamics.

Given the heterogeneous functions that have been ascribed to the theta rhythm, it is possible that it plays a more generalized role in neural processing that is tailored to the structures recruited in each cognitive state^48^. For example, theta could be a mechanism for synaptic modification with experience and for routing and integration^24^ of multimodal information between behaviorally recruited circuits. This could explain the observation of theta across numerous cognitive states and its enhanced recruitment as a function of increased sensory and mnemonic processing^16,21^. Theta’s long period theoretically increases its coding capacity and makes it advantageous for the routing of information within and across distributed networks^22^ overcoming the constraints of conduction velocity and longer synaptic delays^13,14^. This is further supported by the notion that the leading cellular contributors to theta generation in the hippocampus, GABAergic interneurons^15,49^, are highly connected, long-range projecting hub cells orchestrating network synchrony^50^. Additionally, individuals with higher PFC-HC white matter integrity tend to have high slow-frequency power recruitment (delta/theta) and better long-term memory^51^. Finally, intracranial theta burst stimulation of the entorhinal region, where hippocampal/neocortical exchange takes place, has been previously reported to improve memory function in patients with epilepsy^52,53^. Altogether, theta may be a key contributor to routing inputs and integration of multimodal information into synapses, which would play a general role across cognitive domains including, but not limited to episodic memory.

Despite the prominent role of the theta rhythm in primate and rodent studies, it is important to note that not all species utilize theta in episodic memory processes. For example, the hippocampal formation in bats exhibits temporal coding and synchronization in the absence of low-frequency spectral power, suggesting that rhythmicity is not necessary for effective phase coding^54,55^. From an evolutionary standpoint, theta rhythmicity could be one of several possible mechanisms that enable memory formation and retrieval.

In conclusion, our results provide a mechanistic account for hippocampal-neocortical interactions by demonstrating a role for 4-5 Hz-mediated intra and inter-regional dynamics during encoding and retrieval to support accurate memory discrimination. Future studies are needed to identify the molecular and cellular processes underlying the observed role of theta and whether 4-5 Hz codes for memory representations directly or facilitates their exchange.

## Methods

### Experiment paradigm

Electrophysiological recordings were conducted while participants engaged in a hippocampal-dependent pattern separation task (Fig. 1a)^56,57^. The task consisted of an encoding and a retrieval phase. During encoding, subjects were shown 120 images, presented one at a time for 2-seconds. Each stimulus presentation was followed by a question prompt of ‘indoor’ or ‘outdoor’, whose duration was determined by subject response time via a button press. During retrieval, 160 images were presented serially and belonged to one of three categories: repeated images (40) from the encoding phase (repeat), similar but not identical images (80) to those presented during encoding (lure), and new images (40) (new). Following each 1-sec stimulus presentation period, a question prompt of ‘new’ or ‘old’ appeared, to which subjects responded with a button press. A ‘new’ response to a lure image (lure+) reflects successful discrimination between encoding and retrieval images. Conversely, an ‘old’ response to a lure image (lure-) reflects failed discrimination.

Lure images were created by altering a variety of features from the original image (color, rotation, detailed elements in the image), while the new item is an entirely novel image (detailed in Stark et al., 2019). Lure similarity was defined after the creation of the images, using a separate experiment involving healthy participants (n = 116), which tested the probability of falsely recognizing each lure (calling a lure “old”). These ratings were used to rank order lures in terms of their “mnemonic similarity”^58^.

### Participants

Electrophysiological recordings were acquired from eight adult (ages 23-52) patients (3 females) stereotactically implanted with depth electrodes (Integra or Ad-Tech, 5-mm interelectrode spacing) for clinical monitoring (Table 1). Surgical implantation was done to localize the seizure onset zone for possible surgical resection. Electrode placement was determined exclusively by clinical needs and took place at the University of California Irvine Medical Center. The inclusion criterion was concurrent electrode coverage over both hippocampal (HC) and neocortical (NC) areas. Informed consent was obtained from each participant for study inclusion in accordance with the institutional review board of the University of California, Irvine.

### Electrophysiological recording acquisition and data pre-processing

#### Recording and offline filtering

The task was programmed in PsychoPy2 (Version 1.82.01). Participants performed the task on an Apple Macbook Pro placed at a comfortable viewing distance. Participants indicated their responses using an external apple keypad. A photodiode was placed in the corner of the computer screen to capture stimulus presentation onset and offset timestamps with millisecond precision to time-lock them to the neural recordings.

Electrophysiological recordings were acquired using a Nihon Khoden recording system (256 channel amplifier, model JE120A), analog-filtered above 0.01 Hz and digitally sampled at 5000 Hz. Following acquisition, data were pre-processed using customized MATLAB scripts.

Post-acquisition, data were downsampled to 500 Hz. Line noise and harmonics were removed using a 2^nd^ order butterworth band stop infinite impulse response filter, with band stop frequencies centered around (+/−1Hz) 60Hz and 2^nd^-3^rd^ harmonics.

#### Artefact rejection

Artefacts were identified in the time domain for each channel. Time points with values exceeding the mean channel signal +/− 3-4.5 multiples of the standard deviation were indicated. Timepoints near (1-sec before and after) the identified artefacts were also flagged as artefacts. After acquisition of spectral values, flagged artefactual timepoints were removed from the estimates of the mean and standard deviations of spectral power for baseline normalization. Moreover, any trial with a minimum of one artefact time point in the pre-stimulus baseline or post stimulus periods was removed entirely from all analyses.

#### Re-referencing

A Laplacian referencing scheme was implemented, whereby for a given electrode N, the averaged signal was computed over its neighboring electrodes, N-1, and N+1, and subtracted from the signal in electrode N. This has been previously suggested as an optimal scheme for slow frequency oscillatory activity^59^. However, all reported results were reproduced using two additional approaches: 1) No post-acquisition re-referencing and 2) Using a probe-specific white-matter reference electrode.

### Electrode localization and selection

#### Localization

Post implantation CT was co-registered to pre-implantation and post-implantation structural T1-weighted MRI scans for identification of electrode anatomical locations. There were two exceptions; for one patient, a pre-implantation T2-weighted MRI scan was used, and for a second patient, only a post-implantation T1-weighted MRI scan was used for identification.

Overlaid images were visualized in AFNI. An experienced rater (M. S. L.) identified electrodes in neocortical areas which included orbitofrontal (OFC), frontal (FRO), temporal (TEMP), cingulate (CING), insular (INS), and entorhinal/perirhinal (EC/PRC) cortical electrodes as well as hippocampal areas which included CA1, subiculum (SUB), and DG/CA3. OFC labeled electrodes are mutually exclusive from FRO electrodes. The first rater used a previously described^60^ 0.55mm isotropic resolution in-house anatomical template, the Duvernoy atlas^61^ and reported medial temporal lobe (MTL) localization guidelines^62^ for anatomical identification of MTL electrodes. The labeled in-house template was resampled (1mm isotropic) and aligned to each subject’s pre-implantation scans using ANTs Symmetric Normalization^63^. For all other non-MTL electrodes in the neocortex, Freesurfer cortical parcellations^64^ and whole human brain atlas^65^ were used.

Based on the anatomical labels within each subject’s space, the electrode location was determined by identifying the region of interest that encompassed the center of the electrode artefacts. Cases in which electrodes were on the border between ROIs, bordering CSF, or between gray matter and white matter were noted as such and excluded from these analyses. A notable exception from this exclusion criteria are the entorhinal/perirhinal localized electrodes that only bordered white matter or CSF, as electrodes localized in this region are few across all patients.

A second, experienced rater (M.A.Y.) independently localized entorhinal/perirhinal, hippocampal and DG/CA3. Across the two raters there was a 100% concordance for hippocampal sites, 100% concordance for DG/CA3 sites, and 75% concordance for entorhinal/perirhinal sites (those two sites were collapsed during analysis).

#### Electrode selection

The following inclusion criteria were applied for all electrodes (Fig. 1c): 1) Either located in the neocortex or the hippocampus, 2) placed entirely in gray matter (not overlapping with white matter or ventricles) 3) Cue responsive. Neocortical electrodes included: OFC, FRO, TEMP, CING, INS, EC/PRC. Hippocampal electrodes included: SUB, CA1, DG/CA3. Cue responsivity was defined as significant increases in slow frequency power (3-6 Hz) from pre-stimulus baseline (−0.3-0 sec vs. 0-1 sec) in any condition type (repeat+, lure-, lure+, new+), or across pooled trials from all condition types during the retrieval phase. Significant increases were measured using non-parametric, unpaired permutation testing (1000 permutations) between mean pre-stimulus and post-stimulus trial power values for each channel. Contacts from each subject meeting each of the inclusion criteria were projected onto a standard brain model (Montreal Neurological Institute, MNI) and visualized using DSI Studio. Retrieval cue responsive electrodes were used for all analyses (encoding and retrieval power and phase transfer entropy measures).

### Spectral analysis

#### Power extraction and baseline normalization

Analytic Morlet Wavelets were generated in MATLAB using the Wavelet toolbox. There are 4 input parameters to this toolbox: 1) minimum frequency, set to 3, 2) maximum frequency, set to 200, NumVoices, set to 32, and wave center frequency, set to 6/2pi. The toolbox generates ‘scales’ based on the desired frequency range (defined by min to max frequencies), which then get mapped into frequencies. The trial vector, scales vector, and ‘morl’ are inputs to the cwtft function in matlab, which generates the wavelets and power extraction. Wavelets were tested on simulated sinusoids of known spectral properties. Wavelets were used to extract power values in the 3-200 Hz frequency range, which were then converted to decibels (dB). For each trial, instantaneous power values were normalized relative to the channel mean and standard deviation of pre-stimulus power values at the same frequency. For a given trial, the pre-stimulus durations utilized to generate the normalizing distribution were only those preceding the same condition type as the trial to be normalized (condition specific pre-stimulus normalization). Then, for each trial, a z-score value was obtained for each time-frequency point; thereby generating a spectrogram. Baseline normalization was computed separately for the encoding and retrieval phases. Pre-stimulus artefact timepoints were included in the wavelet analysis to maintain the temporal structure of the data but were excluded from the normalizing distribution. A condition - specific baseline was utilized in this study due to condition-specific activity occurring in the response period, which precedes subsequent trials. However, results reported were reproduced with normalization relative to power distributions across the 1) entire recording, and 2) pre-stimulus baseline.

#### Group and individual subject analysis

For condition specific analysis, data were epoched relative to stimulus presentation and subject response timestamps. Z-score normalized spectrograms were averaged across trials within a channel to generate mean z-score normalized spectrograms for each channel. For stimulus onset locked activity, mean spectral power across pooled channels during the 0.2 sec prior to stimulus presentation was removed from the entire power series in the same frequency range. This was done to ensure that power time traces began at a 0 offset at stimulus presentation, which enabled the comparison of post-stimulus power differences between conditions.

For region-based group spectral analysis, channels were pooled across subjects and defined as observations. For region-based individual subject analysis, spectrograms were averaged across channels in each trial, and trials were defined as observations.

#### Identification of significant time-frequency clusters

Significant time-frequency clusters which differed between the lure+ and lure-conditions were identified using cluster-based permutation testing^66^. Briefly, this involved calculating a t-statistic in each time-frequency voxel, between the lure+ and lure-time-frequency z-score normalized matrices (z-maps), thereby generating the observed t-map (2-D matrix). The observed t-map was then compared to a null distribution of t-maps generated over 1000 paired permutations. In each permutation, condition labels were shuffled within a random subset of channels (paired). A p-value for each time-frequency entry was obtained by comparing the observed to the null t-value at the same time-frequency entry, thereby generating a p-map. To correct for multiple comparisons, clusters of contiguous voxels with a *p* <0.5 were identified and compared to the null-distribution cluster size. Observed clusters with sizes larger than the 95th percentile of those from the null distribution were considered significant after correction for multiple comparisons.

### Phase transfer entropy analysis

An information theory metric, phase transfer entropy (PTE), was used to estimate the direction of information transfer in the 4-5 Hz frequency range^19,67^. PTE quantifies the mutual information between the past of signal ‘X’, X(t-*τ*) and the present of signal ‘Y’, Y(t). Mutual information reflects the degree of the reduction in uncertainty of one random variable (distribution of present phase values in region Y), given knowledge of another random variable (distribution of the past phase values in region X). PTE calculation^19^ was performed as described below and normalized between −0.5 and 0.5 as previously reported^67^. PTE was calculated using two methods: first as a time-average measure across the 2-second and 1-second periods following stimulus onsets in encoding and retrieval, respectively. Second, PTE was calculated in moving windows in these same periods (see PTE as a function of time (PTE FOT) below).

Briefly, PTE was calculated using the following formula,

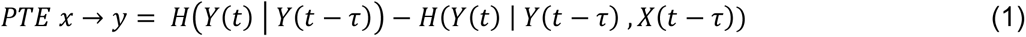

Where *H*(*Y*(*t*) | *Y*(*t* − *τ*)) is the conditional entropy of the distribution of *Y*(*t*) conditioned on *Y*(*t* − *τ*), and *H*(*Y*(*t*) | *Y*(*t* − *τ*), *X*(*t* − *τ*)) is the conditional entropy of the distribution of *Y*(*t*), conditioned on *Y*(*t* − *τ*) and *X*(*t* − *τ*)).

The entropy of a random variable Y is,

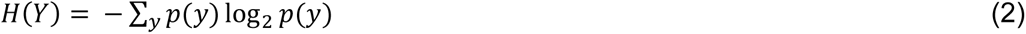

and conditional entropy is calculated by,

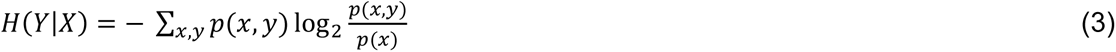

Given these definitions, the terms necessary to compute PTE _X→Y_ are the estimated marginal probability mass function (pmf) of the output past *Y*(*t* − *τ*), joint pmf of the output past and present *Y*(*t*), *Y*(*t* − *τ*), joint pmf of the output and input present *Y*(*t*), *X*(*t*), and joint pmf of the output present, output past and input past *Y*(*t*), *Y*(*t* − *τ*), *X*(*t* − *τ*). Each pmf was estimated using the entirety of the trial data, rather than a single trial. Bin width was defined according to Scott’s choice^68^ and the lag value *τ* was set to 100ms.

PTE _X→Y_ (NC → HC) and PTE _Y→X_ (HC → NC) were calculated separately and normalized as follows:

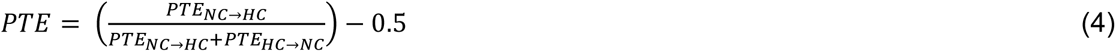

Normalization (dividing by the sum of PTE in both directions) was performed because it is difficult to attribute meaning to the raw unnormalized phase transfer entropy measures; there is no upper bound to entropy values^19^ and there is difficulty in assessing whether values that are close to zero are meaningful^69^. Normalized PTE was then subtracted from 0.5, with a value of 0 denoting bidirectional interactions (or no exchange in the two directions), a value greater and less than zero reflecting higher NC→HC and HC→ NC directional bias, respectively. For all PTE analyses, NC refers to any cue responsive orbitofrontal, frontal, and temporal electrodes.

#### Phase transfer entropy as a function of time (PTE FOT)

PTE as a function of time was calculated as described above, however, in moving 0.5 sec windows with a 10ms step size (encoding and retrieval) and a 50ms step size (encoding).

### Statistical analysis

After identification of the time-frequency cluster which significantly differed between lure+ and lure-across channels in each region, power in this cluster was averaged in each channel, generating a single value. Mean channel cluster power values were used to statistically examine two additional contrasts, lure+ vs. new+ and lure+ vs. repeat+, using paired non-parametric permutations testing; this involved generating a null distribution of differences by randomly shuffling condition labels (in a manner similar to what is was done for the CBPT analysis) and obtaining a p-value by quantifying the number of times a null difference exceeded the true difference, divided by the total number of permutations (n=1000). This was followed by false discovery rate (FDR) correction for multiple comparisons.

Similarly, for assessing statistical differences in PTE between conditions, paired non-parametric permutations testing was performed, as suggested by the initial manuscript that introduced PTE^19^ and later applied by subsequent studies^67,69^. Permutation testing between conditions (or between real and surrogate data) is suggested since PTE values are unbounded^19^. PTE values from NC-HC channel pairs were used as observations. A total of 1000 permutations were performed in all tests to generate null distributions, whereby the condition labels were randomly shuffled in each permutation and a null difference between the two fake conditions was measured. A p-value was again obtained by quantifying the number of times a null difference exceeded the observed true difference.

For all significant results, effects sizes were reported in the form of *η*^2^for parametric tests (repeated measures ANOVA for PTE FOT) or in the form of an effect, defined as the magnitude of the difference between the contrasted group means for non-parametric permutation tests. For the latter, an effect size could not be obtained due to the non-independence between sample observations, limiting the ability to estimate an independent sample standard deviation. More specifically, in using pooled channels (or channel pairs for PTE) as observations, channels coming from the same subject are more correlated thereby decreasing the standard deviation estimate.

In addition, for PTE, chance statistical analysis was performed for retrieval lure+ and encoding →lure+ conditions. This was done by generating a null distribution using the experimental data, by shuffling the phase time series for the sender channel data (NC channels for retrieval and HC channels for encoding) while preserving the temporal structure of the receiving channels. A total of 1000 permutations were performed, whereby in each iteration, this was done for the sender channel of all channel pairs pooled from all subjects, and then averaged across such channel pairs to obtain a mean X → Y null PTE. In total, this yielded 1000 null mean X →Y PTE. Then the true observed value of mean X→Y PTE across all channel pairs using the non-phase shuffled data was examined against the null distribution. A p-value was obtained by quantifying the number of times a null PTE value exceeded that of the observed, divided by 1000.

### Data and Code Availability

All data from this manuscript is available on Open Science Framework (OSF) at the following repository address: https://osf.io/dn9as/. All Code and Statistics files are on GitHub at https://github.com/Yassa-TNL/theta_physiology for open data sharing and to enhance reproducibility.

## Author Contributions

S.G. contributed to design, data processing, data acquisition, data analysis, interpretation, drafting and revision of the paper. M.S.L. contributed to data processing and revision of the paper. L.M., I.S-G, S.V. contributed to data acquisition. L.S. contributed to revision of the paper. P.E.R. contributed to revision of the paper. J.L.L. contributed to design, interpretation, and revision of the paper. M.A.Y. contributed to design, interpretation, and revision of the paper.

## Acknowledgements

We acknowledge contributions to data collection from Dr. Rebecca Stevenson and Dr. Ivan Skelin, as well as the technicians and nurses of the UC Irvine Epilepsy Monitoring Unit. We are especially indebted to the patient volunteers at the UC Irvine Comprehensive Epilepsy Program. We thank Dr. Ken Norman and Dr. Sebastian Michelmann for helpful discussions. Funding for this project was provided by the National Institutes of Health grants R01MH102392 and R01AG053555 (to M.A.Y.), and training grant support T32NS45540 and T32GM008620 (to S.G.) as well as the UC Irvine School of Medicine and the Roneet Carmell Memorial Endowment Fund support (to J.J.L). We would like to acknowledge support from the Uniformed Services University and the Defense Medical Research and Development Program (P.E.R.). The opinions and assertions contained herein are the private opinions of the authors and are not to be construed as official or reflecting the views of the United States Department of Defense or the Henry M. Jackson Foundation for the Advancement of Military Medicine, Inc.

