## Supplementary Figures for "Theta mediated dynamics of human hippocampal-neocortical learning systems in memory formation and retrieval"

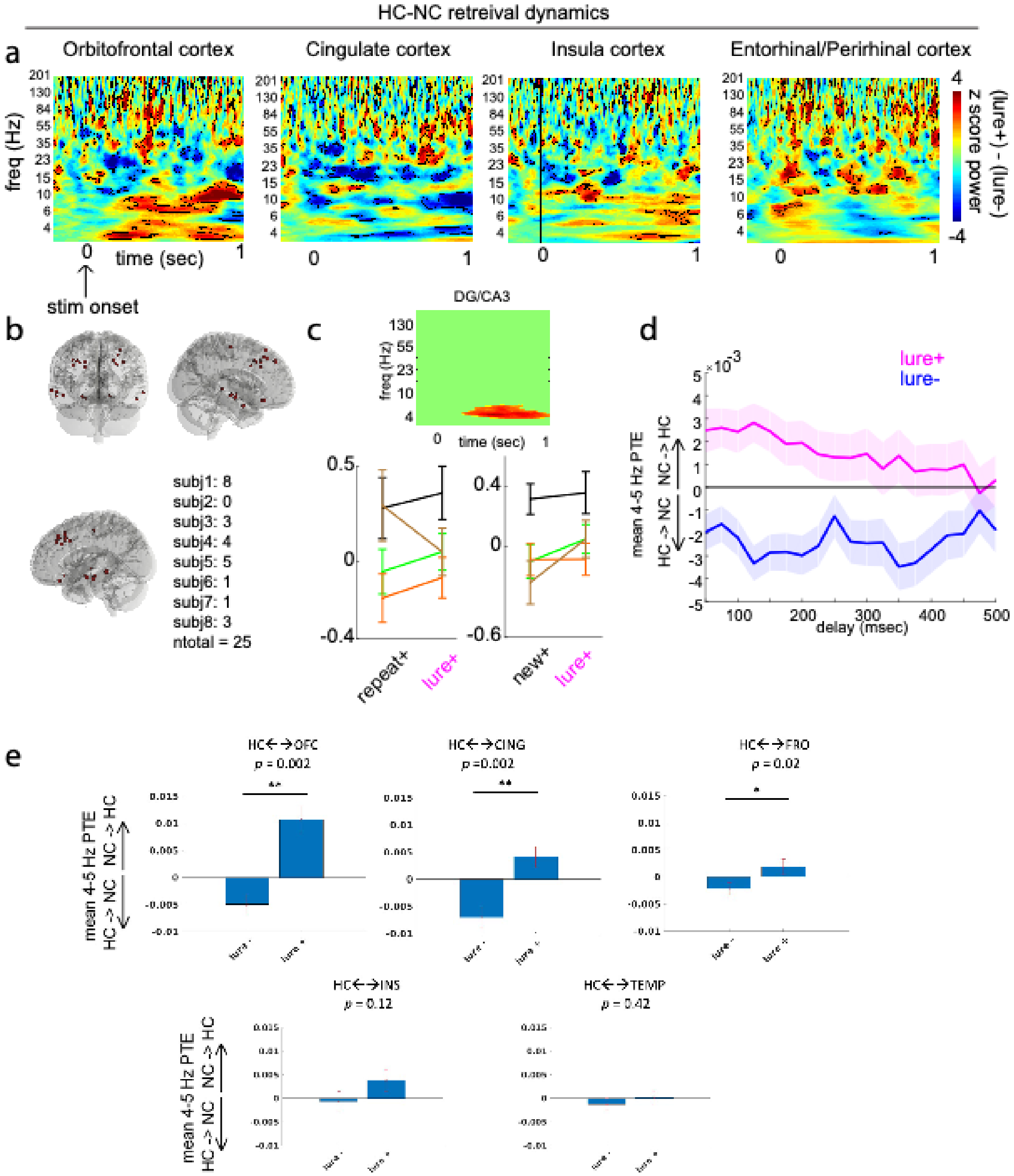


**Supplementary Figure S1. Hippocampal and neocortical dynamics during retrieval. a,** Unthresholded z-score difference map (lure+ - lure-) in orbitofrontal (OFC), cingulate, insular, and entorhinal cortices maps (outlined contiguous voxels, p < 0.05 before multiple comparisons correction). Clusters were not significant after multiple comparisons correction. **b,** Spatial map of OFC, FRO, and TEMP sites, in standardized (MNI) brain space, which showed significantly higher NC theta cluster power in the lure+ compared to the lure- condition (*p* <0.05). Text indicates the number of channels per subject. **c,** Individual subject mean DG/CA3 theta cluster power (cluster shown in right panel) for repeat+ and new+ condition contrasts with lure+. **d,** PTE as a function of different lags (time offset between the NC and HC systems; see methods). Note: a 100 msec lag was used in Fig. 3b. **e,** PTE during the 1-sec post-stimulus period shown separately for each of the NC sites.

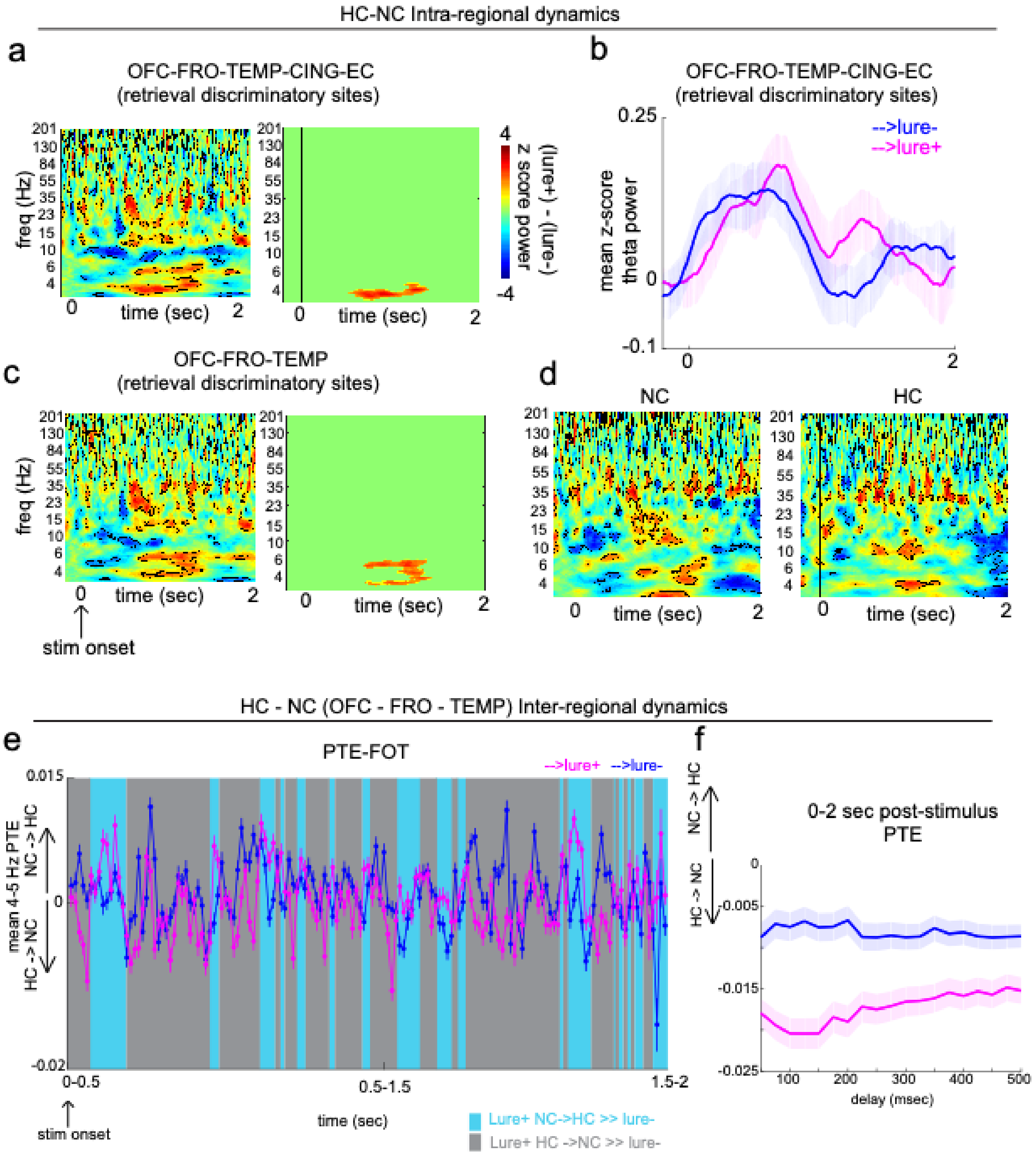


**Supplementary Figure S2. Hippocampal and neocortical intra- and inter-regional dynamics during encoding. a,** Unthresholded (outlined contiguous voxels with p < 0.05, paired permutation testing, 1000 permutations) and thresholded (significant clusters after multiple comparisons correction, p<0.05) z-score difference (→lure+ - →lure-) in all retrieval discriminatory NC sites. **b,** Mean theta (4-5 Hz) power traces across NC discriminatory sites. **c,** Same as in (a), but limiting retrieval discriminatory sites to OFC-FRO-TEMP. **d,** Unthresholded difference maps for all NC and all HC sites. **e,** PTE-FOT relative to stimulus onset, calculated over a 0.5 sec window and a 10ms overlap. Error shades and bars represent the S.E.M across channels (power) or channel pairs (PTE). **f,** PTE as a function of different lags (time offset between the NC and HC systems; see methods). Note: a 100 msec lag was used in Fig. 4d.

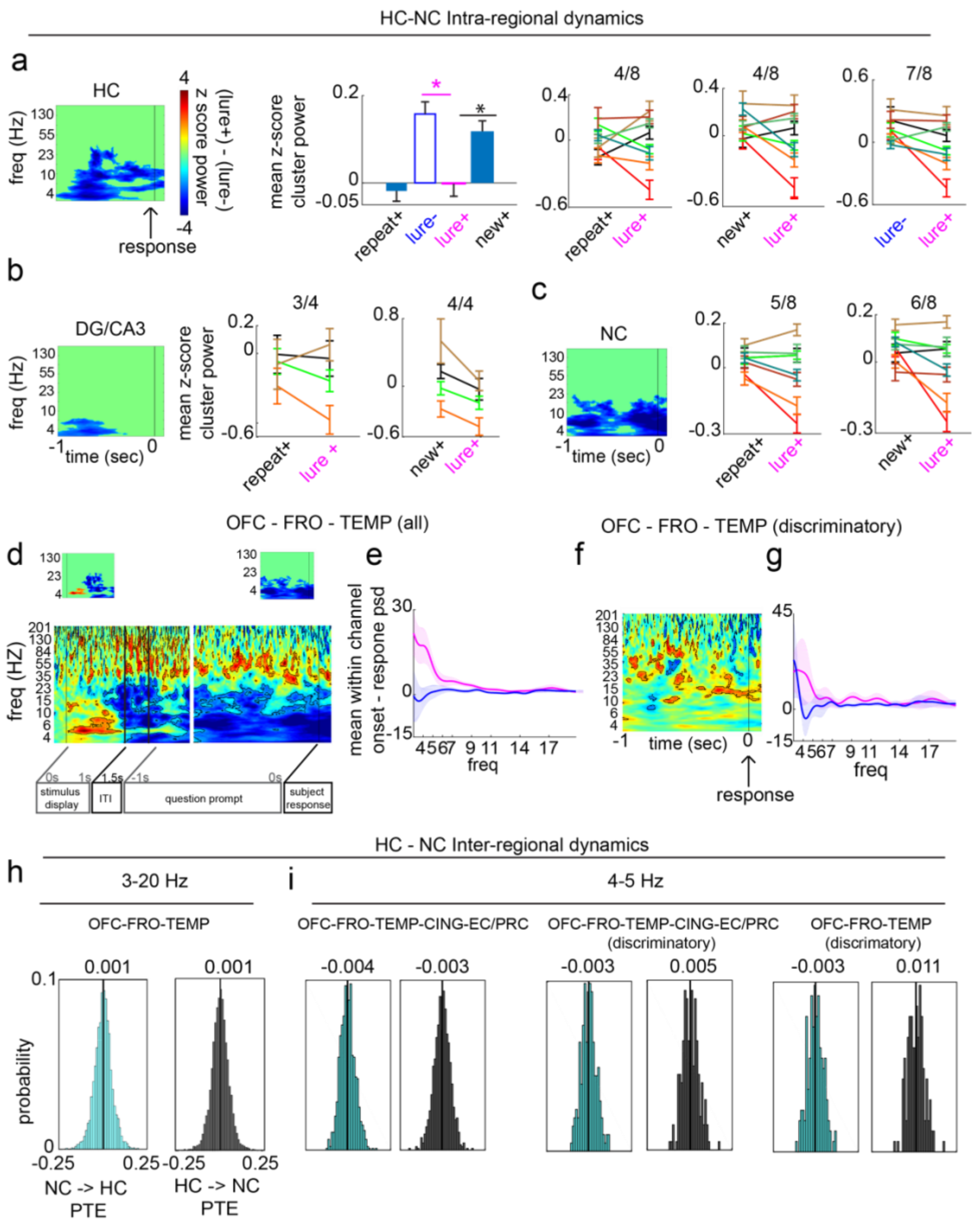


**Supplementary Figure S3. Hippocampal and neocortical intra- and inter-regional dynamics during retrieval response period. a,** HC slow frequency cluster (inset) power, relative to response, was significantly lower in the lure+ compared to the new+ condition ($p$ = 0.002), but did not significantly differ from the repeat+ condition ($p$ = 0.6913, $p_{(FDR)}$ threshold = 0.0279). Effects for lure+ vs. lure- and new+ contrasts = -0.160 and -0.120, respectively. Individual subject mean cluster power for repeat+ versus lure+, new+ vs. lure+, and lure- vs. lure+ contrasts are displayed. **b-c,** Individual subject mean slow frequency cluster power in DG/CA3 (c) and NC (d), relative to response, for repeat+ versus lure+, and new+ vs. lure+ contrasts. **d,** Z-score normalized unthresholded and thresholded (insets) difference maps (lure+ - lure-) locked to stimulus onset (left) and response (right) in NC OFC, FRO, and TEMP sites. **e, g,** Within-channel subtraction of the power spectral density (PSD) between the 1-sec post stimulus onset and 1-sec pre-response period (onset - response PSD) in the indicated sites. **f,** z-score unthresholded difference map locked to response in OFC, FRO, and TEMP discriminatory sites. **h,i,** Probability density function of within channel onset minus response (onset - response) NC–> HC (cyan) and HC→ NC (black) PTE. Distributions are displayed for PTE computed across 3-20Hz in steps of 1-Hz (h), and in the 4-5Hz range (i) using the indicated NC sites. Error shades and bars represent the S.E.M across channels (power) or trials (individual subjects). *$p$ < 0.05, **$p$ < 0.01, ***$p$ < 0.001 indicate significant differences using two-tailed non-parametric paired permutation testing defining channel/channel pairs as observations (1000 permutations).

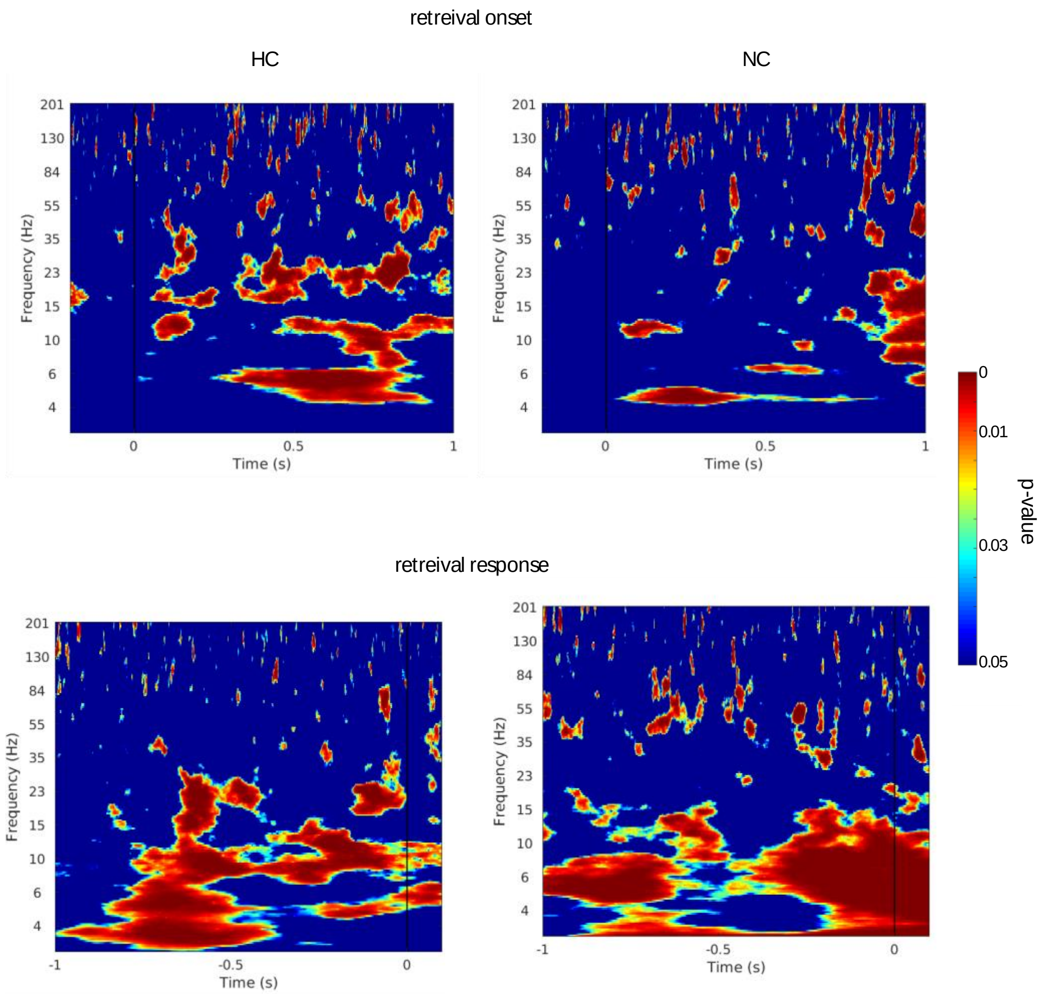


**Supplementary Figure S4. Unthresholded p-value maps.** p-values maps for HC and NC spectrograms (left and right panels, respectively), during the retrieval onset and response task periods (top and bottom panels, respectively). These p-value maps were thresholded again to correct for multiple comparisons based on the size of clusters defined as contiguous entries with a p-value < 0.05. The size of such clusters was compared to the null data cluster size distributions; cluster larger than the 95$^{th}$ percentile of the null were tagged as significant after correction for multiple comparisons, yielding the thresholded maps in Fig 2a bottom left, 2b bottom left and Fig. 4d left top inset and 4a left top inset.

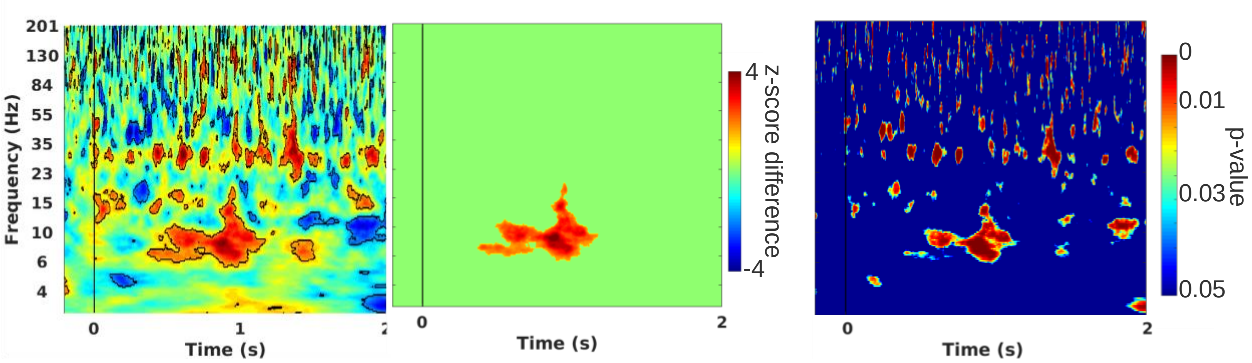


**Supplementary Figure S5. Overall Hippocampal activity during encoding.** Hippocampal sites not limited to retrieval cue responsive contacts (as opposed to all other analyses). (left) Unthresholded (outlined contiguous voxels with $p < 0.05$, paired permutation testing, 1000 permutations) and (middle) thresholded (significant clusters after multiple comparisons correction, $p<0.05$) z-score difference (→lure+ - →lure-) in all hippocampal contacts irrespective of cue responsivity during retrieval (as opposed to figures in Supplementary Figure S2d right panel). Corresponding p-value mapping for the z-score map on the left.

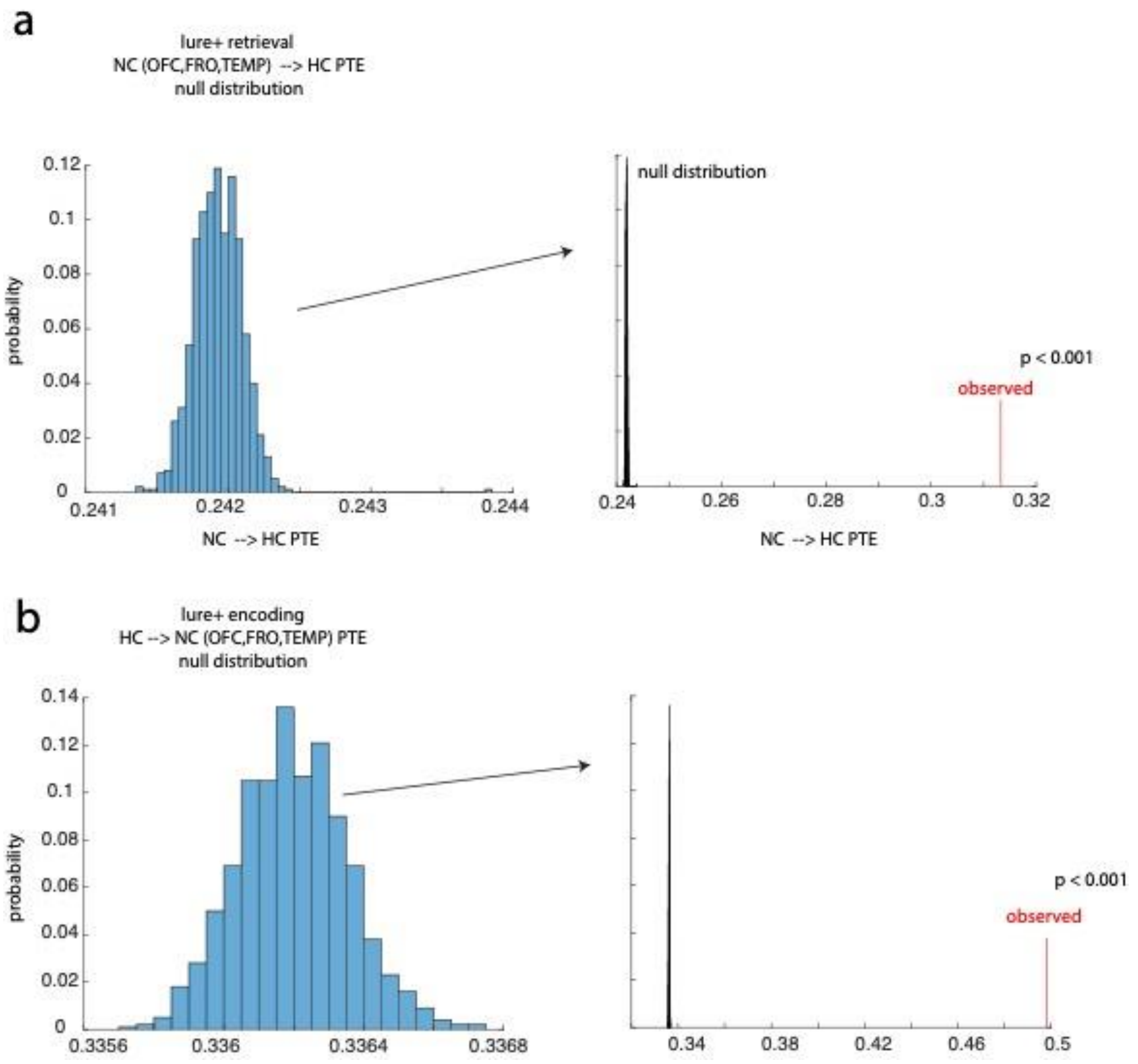


**Supplementary Figure S6. Chance analysis for PTE results. a,** PTE chance analysis for lure+ NC→HC PTE during retrieval. (left) Null distribution was generated by shuffling the phase time series for the NC channel data while preserving HC channel data (n=1000 permutations for each NC-HC channel pair). The probability mass function displayed represents the 1000 samples of the mean null NC→HC PTE across all channel pairs. (right) the observed value shows the mean NC-HC PTE across all channel pairs using the true (non-NC phase shuffled data). Out of 1000 permutations, no single null sample of mean NC->HC PTE exceeded that of the observed value ($p < 0.001$). **b**, (right) same as in (a) but for encoding HC→NC PTE for subsequently discriminated items (→lure+). Shuffling was applied to the HC data (sender channel). (left) Out of 1000 permutations, no single null sample of mean HC->NC PTE exceeded that of the observed value ($p < 0.001$).
